# *De novo* assemblies of high-quality reference transcriptomes identifies Rosaceae-common and *Rosa*-specific encoding genes

**DOI:** 10.1101/199257

**Authors:** Shubin Li, Micai Zhong, Xue Dong, Xiaodong Jiang, Yibo Sun, Li Dezhu, Kaixue Tang, Jin-Yong Hu

**Affiliations:** Flower Research Institute, Yunnan Academy of Agricultural Sciences, National Engineering Research Center for Ornamental Horticulture, Yunnan Flower Breeding Key Lab, Kunming 650205, China.; Group of Plant Molecular Genetics and Adaptation, Key Laboratory for Plant Diversity and Biogeography of East Asia, Kunming Institute of Botany, Chinese Academy of Sciences. Kunming 650201, China.; Germplasm Bank of Wild Species, Kunming Institute of Botany, Chinese Academy of Sciences, Kunming, 650201, China

**Keywords:** reference transcriptome, Rosaceae-common transcripts, *Rosa*-specific transcripts, trinity, RNA-seq, BUSCO, *Rosa*

## Abstract

Roses are important plants for human beings with important economical and biological traits like continuous flowering, flower architecture, color and scent, that current model plants do not feature, thus are becoming ideal models for studying these traits. Due to high heterozygosity of rose genomes likely caused by frequent inter-species hybridization, a high-quality and well-annotated genome for *Rosa* plants is not available yet. Developing genetic and genomic tools with high quality has become necessary for further roses breeding and for disentangling the molecular genetic mechanisms underlying roses domestication. We here generated the high quality and comprehensive reference transcriptomes for *Rosa chinensis* ‘Old Blush’ (OB) and *R. wichuriana* ‘Basyes’ Thornless’ (BT), two roses contrasting at several important traits. These reference transcriptomes showed transcripts N50 above 2000bp. The two species shared about 23310 transcripts (N50 = 2364bp), among which about 8975 orthologs were conserved within genera of *Rosa*. *Rosa* plants shared about 5049 transcripts (Rosaceae-common) with these from *Malus, Prunus, Rubus*, and *Fragaria*. Finally, a pool of 417 transcripts unique to *Rosa* plants (*Rosa*-specific) was identified. These Rosaceae-common and *Rosa*-specific transcripts should facilitate the phylogenetic analysis of Rosaceae plants and investigation of *Rosa*-specific traits. The data reported here should provide the fundamental genomic tools and knowledge critical for understanding the biology and domestication of roses.

## Background

Understanding the molecular mechanisms underlying the adaptation of woody plants to local environmental conditions remains a big challenge in biology due to their long and perennial life history. However, woody plants represent a large proportion of biodiversity on the earth and harbor many different phenological traits that herbaceous plants do not feature (https://www.worldwildlife.org). One such trait is the continuous flowering behavior of modern roses, an important crop for human beings. Guaranteeing a constant supply of raw materials for cut flowers and related products, continuous flowering becomes one of the most important biological and economical traits for roses [1]. Therefore, the genetic control and related gene-regulatory-network for continuous flowering regulation attracts many efforts for many years not only from scientists but also from breeders [1, 2]. Currently, *RoKSN,* the Arabidopsis *TFL1*-like gene in rose, is the only putative target responsible for continuous flowering [3, 4].

Domestication of modern cultivated roses mainly involves hybridization among more than a dozen species [2]. Frequent inter-species crossing/backcrossing and polyploidization of roses has made the classification of roses very difficult though several molecular markers including those from both chloroplast and nuclear genomes have been included in the phylogenetic analyses [5-7]. However, a set of high quality and well-characterized genomic tools/resources are necessary for understanding the biology and domestication of modern roses with more than 30,000 cultivars [8]. Currently, several genetic mapping populations have been developed (see reviews [1, 2]) and determination of rose genomes is underway [9]. On the other hand, due to high-level heterozygosity caused very likely by inter-species crossing and polyploidization, achieving an accurate rose genome seems not so easy. Alternatively, a comprehensive gene expression atlas can be constructed with different tissues at different developmental stages.

The first sets of gene expression atlas were constructed using microarrays containing about 350 (tetraploid *R. hybrida*) [10] and later with about 4800 (*R. chinensis, R. wichuruana*, and *R. hybrida*) [11] selected ESTs. A more comprehensive database containing about 80714 transcript clusters for *R. chinensis* ‘Old Blush’ was constructed from 13 tissues/organs at different developmental stages or under different abiotic and biotic stresses with Illumina and 454 sequencing platforms [12]. Several recent studies have also been done with various *Rosa* species for different purposes [13-16]. Though all these studies have promoted significantly our understanding of the rose biology, the quality of these transcriptomes is normally relatively poor with lower N50 value, completeness, and average length. In this study, we generated a set of high quality reference transcriptomes for two roses by sequencing three tissues at different developmental stages and integrating published datasets. We further identified about 5049 transcripts conserved in Rosaceae plants and 417 transcripts present only in *Rosa* plants. These data should provide fundamental and comparative information for understanding rose biology.

## Data description

### Plant materials and data generation

All samples were collected from a glasshouse located in the Flower Research Institute of Yunnan Academy of Agricultural Sciences (Kunming, Yunnan, China). Leaf materials with about 4cm length (3.5-4.5cm from base of pedicel to leaf tip; at this stage most of the leaflets have just stretched out and become flatten; Figure 1) were collected on 21st Nov, 2015, at when OB was still blooming while BT did not, and on 21st March, 2016, at when both species were setting flower buds (see Figure 2 for workflow). The shoot tip materials with most leaf materials removed were sampled on 21st March, 2016. All the materials were collected with at least six biological replicates at each developmental stage from more than three individual plants of each species.

**Figure 1.**
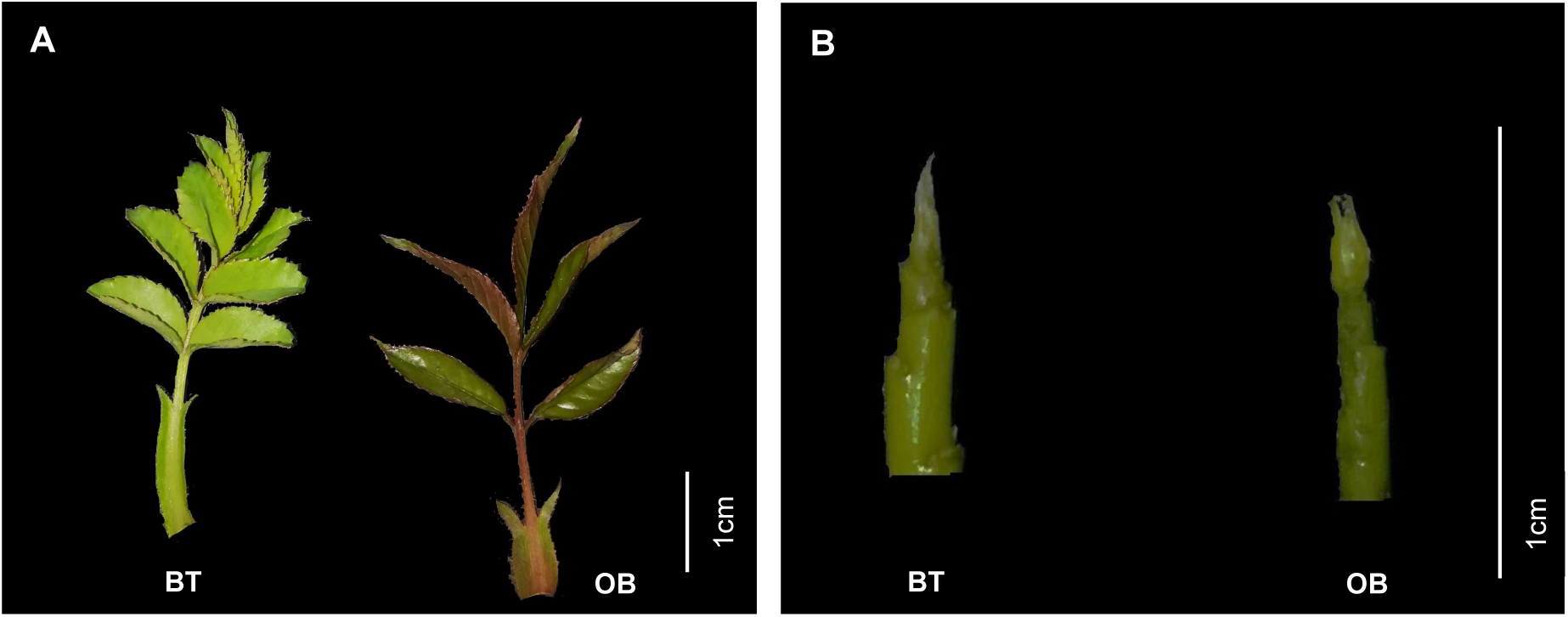
Leaf (A) and shoot (B) materials used for RNA-seq in this study. For each panel, left for *Rosa wichuriana* ‘Basyes’ Thornless’ (BT), and right for *R. chinensis* ‘Old Blush’ (OB). Bars=1cm.

**Figure 2.**
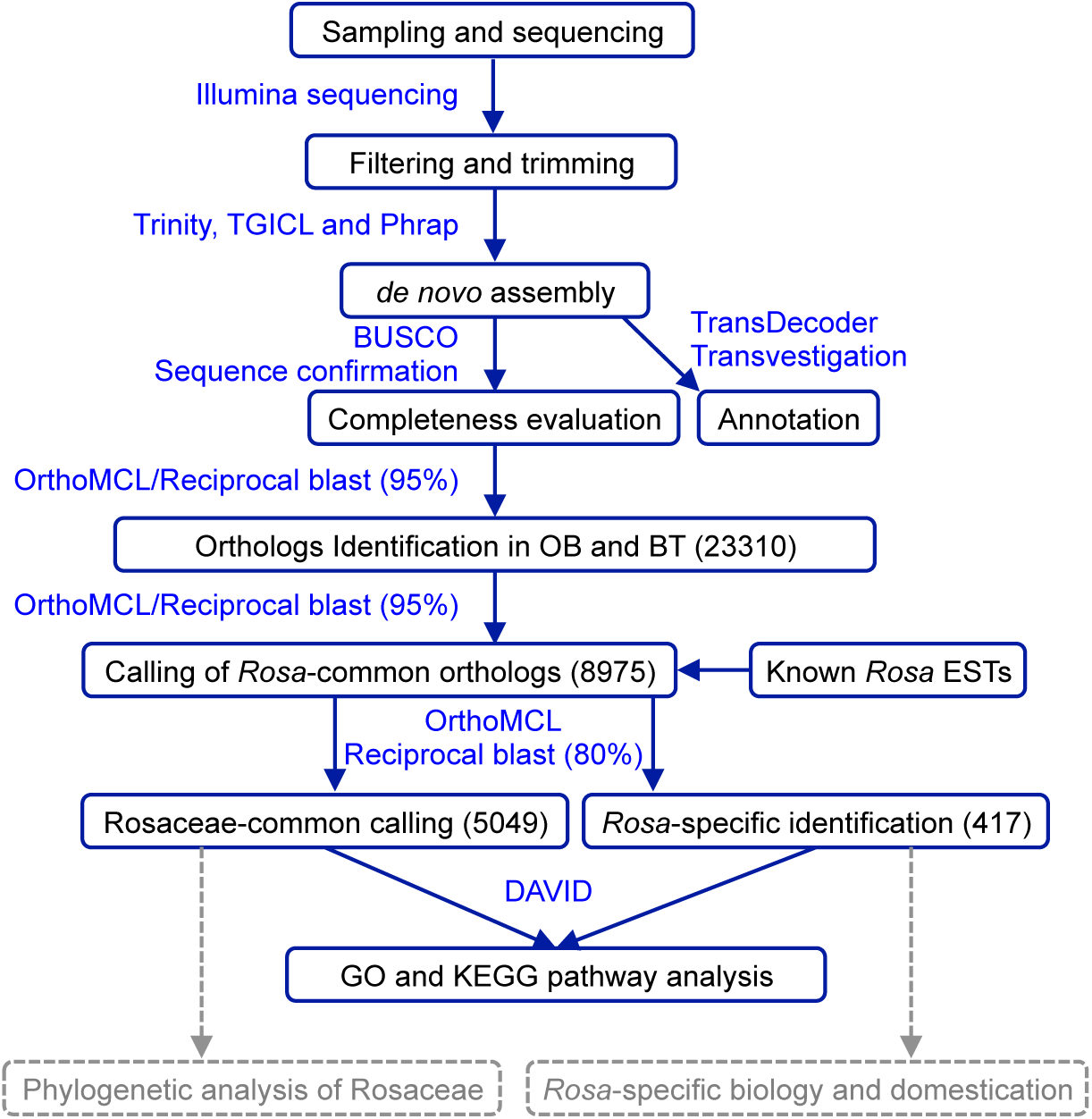
Working flow for assembling of reference trancriptomes and identification of Rosacaeae-common and *Rosa*-specific transcripts. Main steps were shown in boxes with key transcripts numbers given. Major tools used in these analysis were marked in blue. Dashed arrows and boxes indicated the data generated from this study could be explored in these applications.

Total RNA was isolated using the RNAprep Pure Plant Kit (Tiangen, Beijing) and used for mRNA purification with poly-T oligo-attached magnetic beads. Fragmentation was carried out using divalent cations under elevated temperature in an Illumina proprietary fragmentation buffer. Sequencing libraries were generated using the TruSeq RNA Sample Preparation Kit (Illumina, San Diego, CA, USA). First strand cDNA was synthesized using random oligonucleotides and SuperScript II. Second strand cDNA synthesis was subsequently performed using DNA Polymerase I and RNase H. Remaining overhangs were converted into blunt ends via exonuclease/polymerase activities and the enzymes were removed. After adenylation of the 3′ ends of the DNA fragments, Illumina PE adapter oligonucleotides were ligated to prepare for hybridization. To select cDNA fragments of the preferred 380 bp in length, the library fragments were purified using the AMPure XP system (Beckman Coulter, Beverly, CA, USA). DNA fragments with ligated adaptor molecules on both ends were selectively enriched using Illumina PCR Primer Cocktail in a 15 cycle PCR reaction. Products were purified (AMPure XP system) and quantified using the Agilent high sensitivity DNA assay on a Bioanalyzer 2100 system (Agilent). Sequencing was carried out on either Illumina NextSeq 500 or Hiseq2000 platform.

### Data filtering

Approximately 105Gb pair-end data was generated for all samples (about 52.3Gb for BT and 52.8Gb for OB; table 1). The final data volume for OB was about 116.1Gb including the published data. Data information for other species/materials was listed in table 2. The quality of raw reads was assessed and filtered with a custom pipeline using FastQC (V0.10.1; http://www.bioinformatics.babraham.ac.uk/projects/fastqc) and Trimmomatic (V0.36; ILLUMINACLIP:TruSeq3-PE.fa:2:45:10/ LEADING:10/ TRAILING:10/SLIDINGWINDOW:4:25/MINLEN:48) [17]. Adaptor sequences, reads PHRED quality below 92%, and PCR duplicates were all removed with custom perl scripts (https://github.com/ckenkel/annotatingTranscriptomes). Short read archive (SRA) accessions for all data are found in Table 1.

**Table 1.** Summary of sequencing strategies and sequences obtained. Data are sum of three biological replications. ^a^ and ^b^, samples sequenced via Illumina pair-end methods (PE100bp for ^a^ and PE150bp for ^b^); ^c^, data from references (see Table 2).

| Species | Sample | Reads number | Reads bases (nt) | Q20 percent (%) |
| --- | --- | --- | --- | --- |
| *Rosa*<br>*wichuriana*<br>'Basye's<br>Thornless' (BT) | SAM <sup>a</sup> | 156,722,692 | 14,014,983,240 | 95.8 |
|  | leaf_nov <sup>b</sup> | 131,025,452 | 18,922,658,956 | 95.8 |
|  | leaf_mar <sup>b</sup> | 134,110,978 | 19,389,454,378 | 96.1 |
| *Rosa chinensis*<br>'Old Blush' (OB) | SAM <sup>a</sup> | 159,834,774 | 14,385,129,660 | 95.3 |
|  | leaf_nov <sup>b</sup> | 137,399,212 | 19,718,771,811 | 95.8 |
|  | leaf_mar <sup>b</sup> | 130,341,518 | 18,668,608,653 | 96.7 |
|  | all other data <sup>c</sup> | 550,108,308 | 63,356,156,640 | 95.6 |

**Table 2.** Statistics of final assemblies for this study and published data. ^a^, assembly based on data produced from this study; ^b^, assembly based on data from this study and references Yan et al. (2016) and Han et al. (2017); ^c^, conceptual confusion in original text; ^d^, data from Fei lab (http://bioinfo.bti.cornell.edu/cgi-bin/rose_454/index.cgi) with transcript N50 and average length recalculated.

| Assembly components | Contig number | Transcript number | Transcript N50 | GC content (%) | Total assembled bases | Average length (bp) | Data sources |
| --- | --- | --- | --- | --- | --- | --- | --- |
| BT <sup>a</sup> | 86642 | 68612 | 2099 | 46.4 | 92M | 1338 | This study |
| OB <sup>a</sup> | na. | 74975 | 1732 | 44.7 | 88M | 1170 | This study |
| OB <sup>b</sup> | 99456 | 81389 | 2092 | 44.2 | 111M | 1359 | This study |
| OB | na. | 80714 | na | na. | 36M | 444 | Dubios et al. 2012 |
| OB | na. | 68565 | na. | 46.46 | 61M | 887 | Yan et al. 2016 |
| OB | na | 85663 | na | na. | 70M | 814 | Guo et al.2017 |
| OB | 208039 | 111954 | 1997 | 45.8 | 231M <sup>c</sup> | 1111 | Han et al. 2017 |
| *Coreset1*<br>BT vs. OB; 95% identity | na. | 23310 | 2364 | na. | 41M | 1708 | This study |
| *R. multiflora* | 78676 | 61864 | 1907 | 46.03 | 75M | 1216 | Zhang et al. 2016 |
| *R. jacq* cv. gold medal | na. | 80226 | na | na. | 60M | 743 | Gao et al. 2016 |
| *R. roxburghii* | na. | 106590 | na | na. | 37M | 343 | Yan et al. 2015 |
| *R. hybrida* | 93947 | na. | 1589 | na. | na. | na. | Gao et al. 2017 |
| *R. chinensis* 'pallida' | na. | 89614 | Na. | na. | 38M | 428 | Yan et al. 2014 |
| *Rosa* transcriptome <sup>d</sup> | 60944 | na. | 314 | na. | 18M | 302 | Fei lab |
| *Coreset2*<br>All samples; 95% identity | na. | 8975 | 2457 | na. | 20M | 2244 | This study |

### De novo assembly of transcriptomes for OB and BT

Prior to assembly, data for each species was concatenated (SAM, leaf_nov and leaf_mar), and read abundance was normalized to 50X coverage using the *in silico* normalization tool in Trinity [18] to spare assembly time and minimize memory requirements. Assembly for BT was constructed with data generated from this study, while two assemblies were built up first with the newly produced data and later combining with published data (see table 1 and 2). After filtering and normalization the data was about 157 Gb, comprising approximately 1.3 billion normalized read pairs, which were then assembled using optimized parameters (Kmer=2, min_glue=5, SS_lib RF) in Trinity (r2014_07-17) [18]. Transcript expression levels were estimated with RSEM [19] and open reading frames (ORFs) were predicted using Transdecoder (https://github.com/TransDecoder/TransDecoder/wiki). Hmmer3 [20] was used to identify additional ORFs matching Pfam-A domains.

### Quality and completeness evaluation of the transcriptomes

The three assemblies generated from this study had 68612-81389 transcripts with N50 from 1732 to 2099bp and average length from 1170 to 1359 bp, much better than previously published data for these two species (table 2; Figure 3A) [12, 14, 21, 22] and other species/materials [13, 15, 16, 23]. Mean GC content of all assemblies (44.2-46.4%) was comparable to that of published (45.8-46.5%) roses. The completeness of these assemblies was further evaluated with Benchmarking Universal Single-Copy Orthologs (BUSCO) strategy using 1440 near-universal single-copy orthlogs [24]. This analysis revealed a high proportion of complete (C) and single copy (S) from 54.4% to 68.8%, and complete (C) and duplicated (D) from 24.5 to 28.2%. Fragmented (F) and missing (M) BUSCO items occupied about 5.6-21.4% (Figure 3B; Table Figure3B_busco). These results suggest the high quality, completeness, and coverage of these transcriptomes.

**Figure 3.**
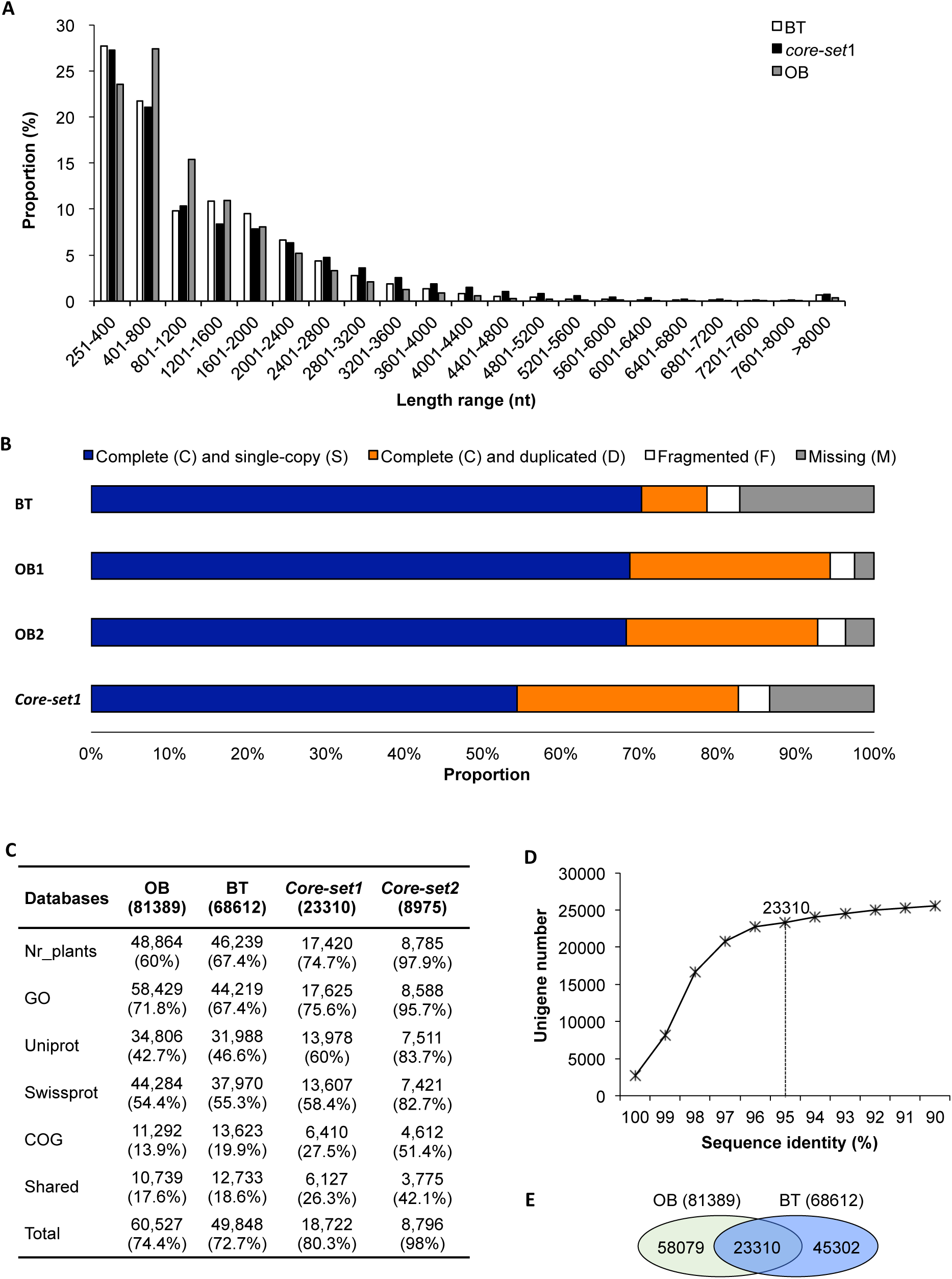
The assembly of high quality transcriptomes for roses. **A.** Length distribution in proportion of assembled unigenes for the two species, *Rosa chinensis* ‘Old Blush’ (OB, bars filled in grey color), and *R. wichuriana* ‘Basyes’ Thornless’ (BT, open bars). Bars filled in black color mark the length distribution of shared transcripts between the two species (*core-set1*; see below and main text). **B.** BUSCO analysis shows the completeness of assemblies and *core-set1*. **C.** Annotation results of the assembled unigenes and *core-set*s for *Rosa*. The *core-set1* is between the two species while *core-set2* is for the unigenes shared at the 95% identity level by the genera of *Rosa* (see Figure 2) based on published and newly collected data from this study. For each category (Nr_plants, GO, Uniprot, Swissprot and COG databases), total unigene counts annotated in different databases besides the proportion (in brackets) are given. Shared and total unigenes annotated by all databases are also given. **D.** Defining the threshold for identifying the conserved RNA element set (*core-set*) between the two species. Unigene number (Y-axis) was plotted against sequence identity (X-axis). Sequence identity at 95% (marked by a vertical dotted line) was chosen for identify the *core-set* unigenes. **E.** Venn diagram shows the results of *core-set1* identification. About 23310 transcripts were identified at the 95% sequence identity level between the two species.

### Functional annotation of transcriptomes

Functional annotation was performed for each of the transcriptomes at the peptide level using a custom pipeline that defines protein products and assigns transcript names. Predicted proteins/peptides were analyzed using InterProScan5 [25], which searched all available databases including Gene Ontology (GO:201605). BLASTp analysis was performed with the UniProt, SwissProt database (downloaded May 2016). The resulting .*gff3* and .*tbl* files were further annotated with functional descriptors in Transvestigator (http://doi.org/10.5281/zenodo.10471).

With available databases explored, more than 72% of the transcripts were annotated for both transcriptomes of OB and BT. The proportion of shared transcripts with annotation in all five databases was about 18% for the two transcriptomes. Detailed annotation information was included in Figure 3C.

### Calling of the <u>c</u>onserved <u>o</u>rthologous transcript <u>e</u>lements <u>set</u> between OB and BT (coreset1)

To identify the transcripts shared and facilitate the gene expression comparison between the two *Rosa* species, we identified the conserved orthologous transcript elements set between OB and BT using orthoMCL [26] and an optimized reciprocal blast method [27] with a sequence identity at 95% (Figure 3D). This analysis identified 23310 transcripts shared by the two species, while OB and BT specific were 58079 and 45302 (Figure 3E). Interestingly, *coreset1* showed more than 80% of the transcripts with annotation (Figure 3C). BUSCO analysis showed that *coreset1* transcripts had quite high completeness (Figure 3B). It’s worthy to note that the total number of *coreset1* could be higher as we used 95% sequence identity as the cutoff of conservation between these two species.

### Identification of coreset2 for Rosa

To screen for conserved transcripts within *Rosa*, we compared *coreset1* to other published RNA-seq data for *R. multiflora, R. roxburghii*, etc (see Table 2 for the data used in this analysis) as no published genome sequences and gene models available. With the sequence identity set at 95% as for *coreset1*, we detected about 8975 transcripts (N50 at 2457bp and mean length at 2244bp) shared for all the plants included (*coreset2*; Table 2). A BUSCO test for transcripts completeness revealed a relative high proportion of fragmented (2.7%) and missing (19.5%) data (Figure1-ST5_busco).

### Identification and characterization of Rosaceae-common and Rosa-specific transcripts

We further compared the *coreset2* transcripts to known CDS for *Malus domestica* (v3.0.a1), *Prunus avium* (v1.0.a1), *Rubus occidentalis* (v1.0.a1), *Fragaria ananassa* (v1.0_FAN_r1.1) (ftp://ftp.bioinfo.wsu.edu/species/) in order to find out the transcripts present in all Rosaceae and only in *Rosa* plants. Reciprocal blast analysis revealed that the total counts shared by all Rosaceae plants varied following sequence identity from 100% to 70% while the slope became saturation at about 85% (Fig2tableSxxx-Rosaceae-common). We then set the cutoff at 85% and detected about 5049 transcripts among *coreset2* were shared by all Rosaceae plants (Rosaceae-common), while only about 417 transcripts present in *Rosa* (*Rosa*-specific; Figure 4A; Fig2tableSxxx-Rosaceae-common; Fig2tableSxxxx-Rosa-specific417x). As the *coreset2* transcripts were used to blast against the CDS from other plants, this analysis did not detect other genera-specific transcripts/genes.

**Figure 4.**
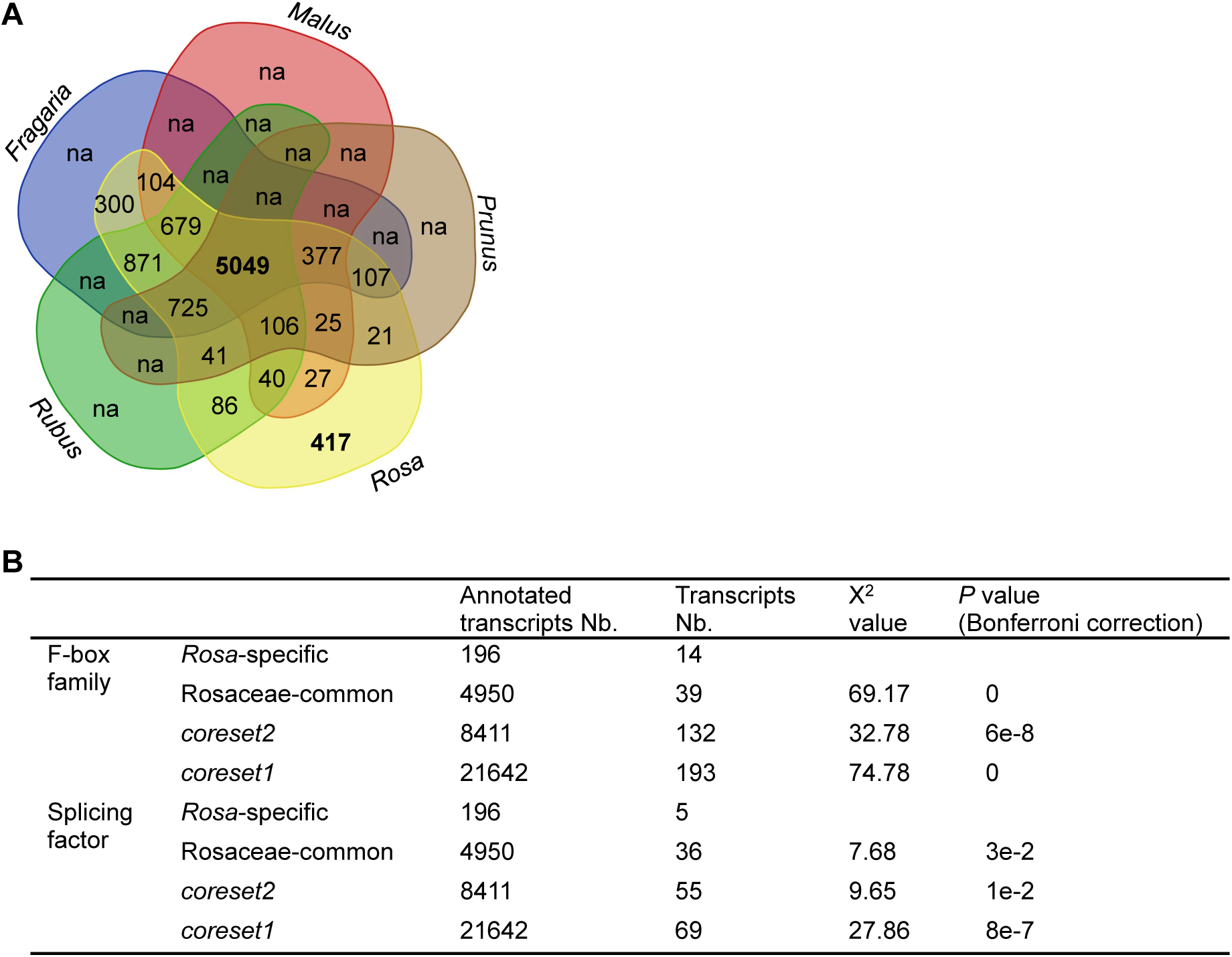
Identification and characterization of Rosaceae-common and *Rosa*-specific transcripts. **A.** Venn diagram shows the Rosaceae-common and *Rosa*-specific transcripts. Note that, except *Rosa*, transcripts specific for other genera were not identified (marked with na). **B.** F-box and splicing factors were significantly enriched in *Rosa*-specific transcripts. X^2^ tests were performed online (http://www.quantpsy.org/chisq/chisq.htm) by comparing the *Rosa*-specific transcripts number with those from Rosaceae-common, *coreset1* and *coreset2* genes. *P* values were corrected with Bonferroni correction.

Most of the Rosaceae-common transcripts were annotated (4950 or about 98%). In these transcripts, there was no significant enrichment of special GO items comparing to *Arabidopsis*, while several GO items related to metabolism (like sugar/carbon/pyruvate, etc) could be detected highly enriched comparing to strawberry genome (Figure Sx). Within the 417 *Rosa*-specific molecules, about 196 transcripts were annotated (in which 50 were uncharacterized and 15 hypothetical) but without significant enrichment of any GO item (Fig2tableSxxxx-Rosa-specific417x). Among these transcripts with functional annotation, four transcripts were related to glycan biosynthesis/metabolism and proteoglycans, while transcripts encoding for F-box family members (14x) and splicing factors (5x) were significantly enriched (Fig. 4B; Fig2tableSxxxx-Rosa-specific417x). F-box proteins represent one of the largest super-families in plants. Splicing factors are important regulators of messenger RNA diversity. Interestingly, some F-box proteins are carbohydrate-binding and presumably targeting glycoproteins for proteasome degradation [28]. Both F-box proteins and splicing factors have been confirmed to play essential roles in almost every aspect of plant growth, development, and adaptation to environmental stresses. It will be then very interesting to evaluate the function of these transcripts/proteins in roses.

## Discussion

As one of the most important horticultural plants, rose has its special biology. Continuous flowering, fragrance, flower shape, thorn and many traits not presenting in *Populus* and other woody model plants could be found in roses, hence roses are now becoming a model woody species for understanding the molecular mechanisms regulating these traits [1]. Interestingly, the breeding of modern roses often involves frequent hybridization and polyploidization among species, which often feature stronger diseases resistance and cold resistance, better fragrance and lack of prickles [29-31]. On the other hand, tracing the processes and history of modern roses domestication and breeding remain a challenge as inter-species crossing and polyploidization was frequently observed [2]. Since a high quality genome assembly for roses is not available yet [9], identification of *Rosa*-specific genes/transcripts might provide a tool for this purpose.

In this report, we produced reference transcriptomes for *R. chinensis* ‘Old Blush’ (OB) and *R. wichuriana* ‘Bayes’ Thornless’ (BT) with transcripts N50 above 2kb and mean length about 1.2kb. Via incorporating published data for OB, we even generated a better assembly with mean transcript length longer than 1.3kb. We identified 23310 conserved orthologous transcripts (*coreset1*) between OB and BT with BUSCO assay confirming the high level of completeness with these assemblies. As *coreset1* transcripts were based on a very high level of sequence identity (95%), they could directly be used to evaluate the differential expression of orthologous genes between species.

Later, these assemblies were explored to identify about 8975 transcripts shared by *Rosa* plants. Finally we detected 5049 transcripts shared by all the Rosaceae plants mentioned in this study, and about 417 transcripts only present in the genera of *Rosa*. The Rosaceae-common dataset contains about 572 single-copy transcripts. These single-copy transcripts could be directly used to clarify the Rosaceae phylogenetic relationship, a challenge likely caused by frequent hybridization, rapid radiation, polyploidization and domestication [32-36].

In contrast to the Rosaceae-common transcripts, the *Rosa*-specific transcripts might be explored to investigate traits specific for roses. Although partial of the domestication processes had been documented [2], the evolutionary history and molecular mechanisms controlling traits special for roses are still not clear [1]. More than half of the 417 transcripts are uncharacterized or without known GO annotation, hence might be related to the phenotypes that have not been characterized in other species. Therefore, these transcripts can be explored to trace trait domestication history in roses.

In summary, these assemblies not only provide much better quality transcriptomes for roses, but also help us pin out the transcripts making roses special.

## Abbreviations

BUSCO: bench-marking universal single-copy orthologs;
GO: gene ontology;
ORF: open reading frame;
SRA: short read archive.

## Author’s contributions

J.H. and X.D. designed and guided the research together with K.T., D.L., and S.L., X.D. S.L., X.J, and Y.S. performed experiments and helped in data analysis. M.Z. and X.D. analyzed data. J.H. and M.Z. wrote the manuscript with the corrections from K.T. All authors have read and approved the final manuscript.

## Acknowledgements

We thank Ms. Dongmin Jin, Dr. Fang Cheng, Ms. Shulan Chen for assistance in experiments. We are grateful to Prof. Chenjun Zhang and his colleagues for accessing to high-performance computers.

## Availability of supporting data

additional file 1: Procedure fig

additional file 2: assembly length distribution fig

additional file 3: statistic of different assembly strategy

additional file 4: ortholog variation with different cutoff

additional file 5: the comparetion between orthoMCL method and reciprocal blast method

additional file 6: BUSCO raw data

additional file 7: the rosaceae CDS data information

additional file 8: the go term clustering for coreset2

additional file 9: the venn fig of the existed EST data base and coreset2

additional file 10: phylogeny tree for the 572 ortholog in rosaceae (none)

additional file 11: the rosa specific transcripts function annotation and enrichment analysis

